# The Amazon River microbiome, a story of humic carbon

**DOI:** 10.1101/2021.07.21.453257

**Authors:** François-Étienne Sylvain, Sidki Bouslama, Aleicia Holland, Nicolas Leroux, Pierre-Luc Mercier, Adalberto Luis Val, Nicolas Derome

**Author notes:** **Corresponding author**: François-Étienne Sylvain. **Competing interests**: The authors declare no competing interests.

## Abstract

The Amazon River basin sustains dramatic hydrochemical gradients defined by three water types: white, clear and black waters. Black waters contain important loads of allochthonous humic dissolved organic carbon (DOC), mostly coming from bacteria-mediated lignin degradation, a process that remains understudied. Here, we identified the main bacterial taxa and functions associated with contrasting Amazonian water types, and shed light on their potential implication in the lignin degradation process. We performed an extensive field bacterioplankton sampling campaign from the three Amazonian water types, and combined our observations to a meta-analysis of 90 Amazonian basin shotgun metagenomes used to build a tailored functional inference database. We showed that the overall quality of DOC is a major driver of bacterioplankton structure, transcriptional activity and functional repertory. We also showed that among the taxa mostly associated to differences between water types, *Polynucleobacter sinensis* particularly stood out, as its abundance and transcriptional activity was strongly correlated to black water environments, and specially to humic DOC concentration. Screening the reference genome of this bacteria, we found genes coding for enzymes implicated in all the main lignin degradation steps, suggesting that this bacteria may play key roles in the carbon cycle processes within the Amazon basin.

## Introduction

The Amazon basin occupies almost 38% of continental South America [1] and holds 17% of the planet’s freshwater [2]. Its discharge carries a significant load of terrestrially-derived nutrients to the ocean, which have global consequences on marine primary productivity and global carbon sequestration [3, 4]. The Amazon River basin sustains dramatic hydrochemical and ecological gradients that impose physiological constraints upon its aquatic communities [5–10]. Its three major tributaries, the Rio Solimões, Rio Negro, and Rio Tapajos, represent distinct water ‘types’ or ‘colors’, which harbor contrasting physicochemical profiles [11]. The white water from the Rio Solimões has an Andean origin, is eutrophic (nutrient- and ion-rich), turbid, and has a circumneutral pH [11–14]. The crystalline ‘clear water’ from the Rio Tapajos has a circumneutral pH, low conductivity, and a reduced amount of suspended material associated to its pre-Cambrian origin draining the Brazilian shield. Lastly, the black water of the Rio Negro stems from the craton born drainage of the Guyana shield [15] and largely contrasts with the aforementioned tributaries: it is oligotrophic (nutrient- and ion-poor) and contains a high quantity of dissolved organic carbon (DOC) - typically 8-12 mg C L^−1^ [10]. DOC from the Rio Negro has a distinctive allochthonous origin, in comparison with Rio Solimões or Rio Tapajos DOC [16]. Terrestrial DOC in the Rio Negro is characterized by an enriched load of humic acids – coming from lignin degradation processes - which acidify the entire Rio Negro watershed (pH 2.8 – 5), making it a challenging aquatic environment for most aquatic species [17].

Overall, the Amazon basin has very low rates of phytoplankton production [18], suggesting that terrestrial DOC is an important carbon source for bacterial growth [19, 20]. This is especially true in the Rio Negro, where most of the DOC comes from plant decomposition, released in the water following the seasonal flooding of a significant part of the forest [11, 21], which provides a concentrated input of lignin-derived components in the environment [22]. Lignin is naturally recalcitrant to degradation [23, 24], since its role is to prevent microbial enzymes from degrading labile cell-wall polysaccharides [25]. However, heterotrophic microbes are able to degrade up to ~ 55% of the lignin produced by the rainforest [26, 27]. The first step of bacterial lignin degradation is the oxidation of lignin-derived compounds, producing a complex mixture of aromatic compounds, which represent the humic fraction of the DOC detected in the Rio Negro’s black water [21, 28].

Despite its relevance for global-scale elemental cycling and primary production processes, there is a limited understanding of the taxonomical and functional structure of the Amazon River microbiome [20]. Also, the transcription profile of taxa involved in Amazonian black water lignin degradation processes has yet to be investigated. In this study, we aimed (1) to identify the most important environmental variables shaping the Amazonian bacterioplankton community structural, transcriptional, and functional profiles; and (2) to better understand the interaction between humic DOC and microbial consortia, by highlighting the taxa likely involved in the lignin degradation process and the potential pathways of degradation they possess.

## Methods

### 1. Sampling and processing

Water samples were collected from 15 sites in the Brazilian Amazon basin between September-December 2018 and 2019. The 15 sites were distributed over an area of >300,000 km^2^ along the Rio Negro, Rio Solimões and Rio Tapajos watersheds, the three major tributaries of the Amazon River [11]. GPS coordinates and a map of all sites are found in Table 1 and Fig. 1, respectively. The 15 sites include five black water, seven white water, and three clear water sites. Six replicate water samples were collected per site. Surface water samples were taken at a depth of 30 cm in 2 L Nalgene™ (Thermo Fisher Scientific, Waltham (MA) USA) bottles. Filtration was performed as in [29] through 22 μm-pore size polyethersulfone Sterivex™ filters (cat #SVGP01050, Millipore, Burlington (MA), USA) less than 30 minutes after collection. Filters were stored in 2 mL of NAP conservation buffer [30, 31] immediately upon collection, and then stored in −80°C until processing. Before DNA/RNA extraction, Sterivex™ filter casings were opened and processed according to [30] using sterile instruments, and filter membranes were stored in TRIzol™ (cat #15596026, Thermo Fisher Scientific). DNA and RNA extractions were performed according to the manufacturer’s instructions of TRIzol™ without modification. Four blank controls (sterile filters stored in the NAP buffer) were also processed identically to all samples for DNA/RNA extractions.

**Table 1:** Site identification, geographical coordinates, dissolves organic carbon (DOC) quantity and quality characterization. “DOC conc.” means DOC concentration in mg L^−1^. SAC340 and SUVA254 are the specific absorbance coefficients index of relative DOC aromaticity (higher the values more aromatic is the DOC). Abs254/365 is the index of molecular weight: the lower the value the higher molecular weight is the DOC is.

**Figure 1:**
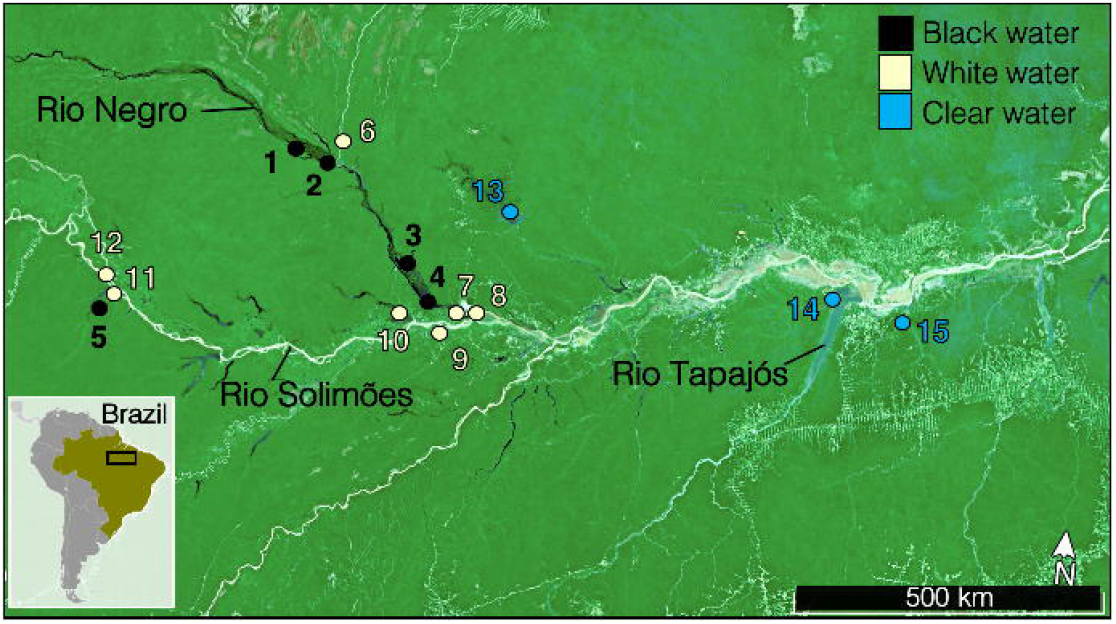
Map of the 15 sampling sites distributed throughout the Brazilian Amazon basin. The color of the dots represent the water type (black, white of clear).

### 2. Environmental variables

A total of 34 environmental variables were measured in this study (see Table 1, Suppl. Table 1, 2, 3). Temperature (°C), conductivity (μS), pH and dissolved oxygen (%) were measured directly on site using a YSI professional plus series multimeter (YSI Inc/Xylem Inc, Yellow Springs (OH), USA). The concentration of DOC, dissolved metals, nutrients, free ions and primary production (i.e. chlorophyll a) were measured at the laboratory, according to the techniques described in details in Suppl. mat. DOC quality was measured using Fluorescence excitation emission (FEEM) scans and modelled using parallel factor analysis (PARAFAC) (see Suppl. mat.).

### 3. Data availability

The datasets generated and analysed during the current study can be found in the Sequence Read Archive (SRA) repository, BioProjectIDs: PRJNA736442 and PRJNA736450. Accession numbers of the 90 metagenomes used to build the custom database are in Suppl. mat. The scripts used for the DNA/RNA sequence analysis, as well as all related input files including all metadata are freely available on the Open Science Network platform (URL: https://osf.io/dz6vf/).

### 4. 16S sequence analysis

Microbial community structure and expression were assessed using a 16S rRNA approach conducted on DNA and RNA extracts. Retrotranscription of the RNA extractions was done using the qScript cDNA synthesis kit (cat #95048-100) from QuantaBio (Beverly (MA), USA) according to the manufacturer’s instructions. Then, the fragment V3-V4 (~500 bp) of the 16S rRNA gene was amplified in DNA and cDNA extracts by PCR using the forward primer 347F (5’-GGAGGCAGCAGTRRGGAAT-3’) and the reverse primer 803R (5’-CTACCRGGGTATCTAATCC-3’) [32]. The PCRs were performed according to the manufacturer’s instructions of the QIAGEN® Multiplex PCR kit (cat #206143, Hilden, Germany) using an annealing temperature of 60°C and 30 amplification cycles. Amplified DNA was purified with AMPure beads (cat #A63880, Beckman Coulter, Pasadena (CA), USA), according to the manufacturer’s instructions, to eliminate primers, dimers, proteins, and phenols. Post-PCR DNA concentration and quality were assessed on a Qubit™ instrument (Thermo Fisher Scientific) and by electrophoresis on 2% agarose gels. After purification, multiplex sequencing was performed on Illumina MiSeq by the Plateforme d’Analyses Génomiques at the Institut de Biologie Intégrative et des Systèmes of Université Laval.

After sequencing, 24,341,734sequences were obtained (mean of 135,232 sequences per sample). DADA2 [33] was used for amplicon sequence variant (ASV) picking. Details on 16S sequence processing including ASV picking and decontamination using control samples is included in Suppl. mat. An analysis of Shannon diversity according to sampling depth for each sample can be found in Suppl. Fig. 1 and 2. ASV tables, metadata files and taxonomy information were incorporated into *phyloseq* objects [34] before downstream analyzes.

### 5. Shotgun metagenome functional database

We built a reference metagenomic database to infer the functional profile of the microbial communities previously characterised using the 16S rRNA approach. The reference database was built from 90 metagenomes from the Amazon River, sampled in previous studies [20, 35–38]. Details on the shotgun metagenomic database construction can be found in Suppl. mat.

### 6. Statistical analysis

#### 6.1 Objective #1: Identify the environmental variables shaping the structural, transcriptional and functional profiles of Amazonian bacterioplankton

First, we aimed to understand to what extent environmental conditions drive the phylogenetic structure, the transcriptional profile and the functional repertory of Amazonian bacterioplankton communities. To do so, we computed Non-metric multidimensional scaling (NMDS) ordinations based on unweighted Unifrac distances [39] between communities (Fig. 2). Biological replicates from every site were merged before this analysis for ease of viewing in the NMDS plot. All the 34 environmental parameters were fitted to the ordinations using *envfit* from vegan [40], however, only the 10 parameters with the strongest R^2^ values were kept for plotting. Ellipses showing a 95% confidence interval were plotted with *ordiellipse* [40]. Permutational Analyses of Variance (PERMANOVA) were performed with *adonis* [40] to test the presence of significant difference between water types.

**Figure 2:**
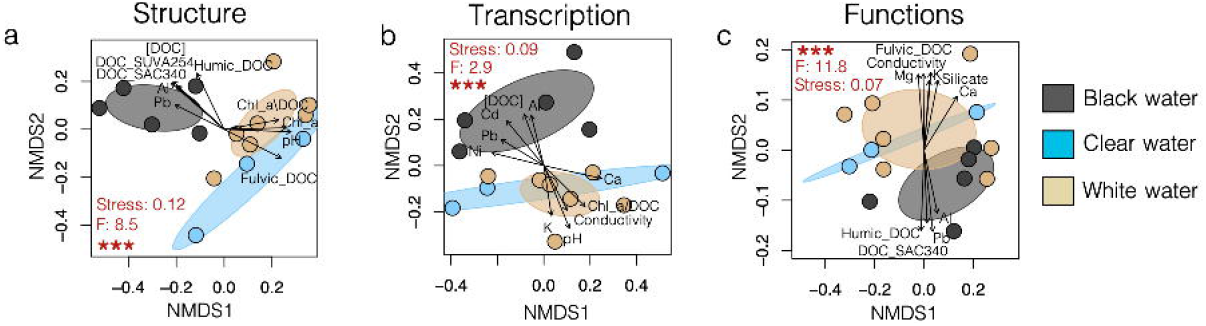
The structure, transcription and functional profiles of bacterioplankton significantly cluster according to water type. NMDS ordination plots of sampling sites according to the (a) bacterioplankton taxonomic structure, (b) transcriptional profile and (c) functional repertory. Samples (n = 90) were merged per sampling site and were colored according to their water type. P and F values from the PERMANOVA tests and NMDS stress values are in red; *** means p-value < 0.001. Only the 10 environmental parameters showing the strongest association to microbiome data (via *envfit*) were represented in each plot for ease of viewing.

Then, we aimed to identify which taxa were associated with the variations in environmental parameters between water types. Although white and clear waters have different origins [11], we decided to merge them together for this analysis, since they show similar environmental parameter profiles in comparison to black water (Table 1, Suppl. Table 1, 2, 3, Suppl. Fig.3). Also, further analyzes focusing on DOC show a unique humic DOC signature in black water in comparison to the other water types. Thus, we were mostly interested in highlighting taxa associated with black water environments rather than to the other water types. We used a machine-learning approach to identify the taxa that best discriminated black versus white/clear waters. To do so, we implemented Breiman's Random Forest algorithm for classification [41] using *randomForest* [42] with ntree = 100. This algorithm splits data in a training and a test set - the training set is used to construct consensus trees of classification via bootstrapping, and the test set (≈37% of the samples) is then used to estimate the node error rates in the trees of classification (i.e. the out-of-bag estimate of error). We isolated the 40 ASVs responsible for the most important mean decrease in GINI coefficient (measure of node purity) with significant p-values following Bonferroni correction. These ASVs comprised the 40 taxa that best discriminated black versus white/clear waters (with the lowest classification error rate) in the random forest tests. The abundance or the transcriptional activity of these taxa in all samples was represented on heatmaps (Fig. 3) along the strength of their GINI coefficient decrease score.

**Figure 3:**
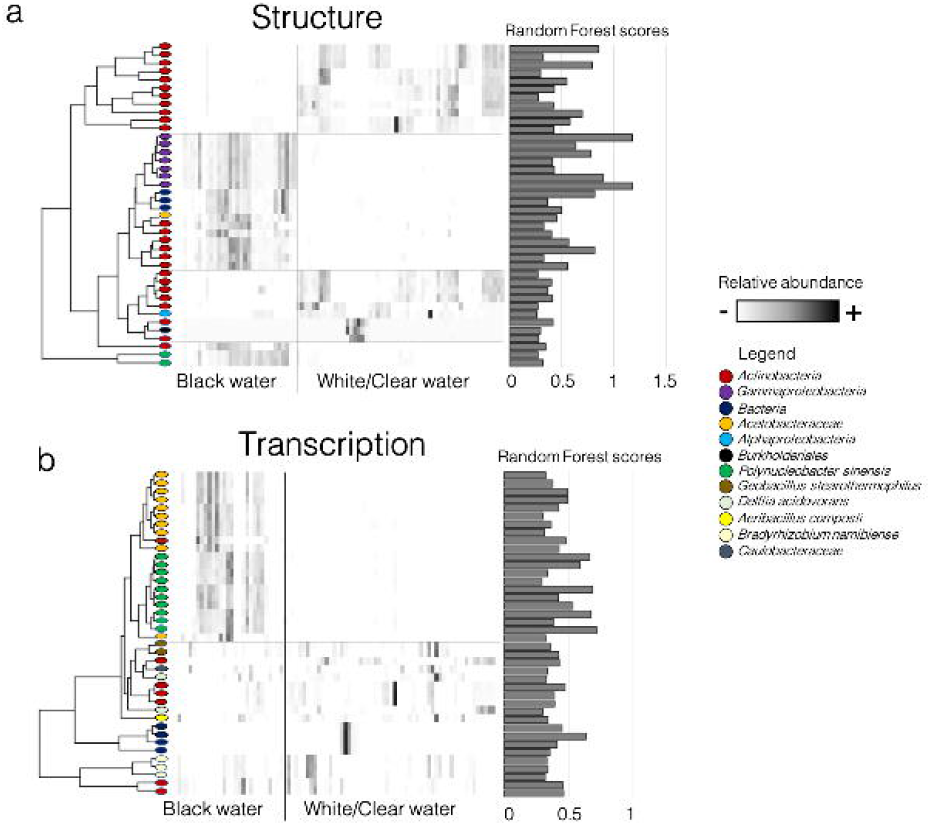
Random-Forest (RF) machine-learning analysis identifies the 40 taxa showing the most important differentiation between black and white/clear water types. Results are shown in heatmaps for the taxonomical structure (a) and the transcriptional profile (b) datasets. Heatmap columns represent samples and rows are different ASVs. The colored shape at the left of each row corresponds to the ASV taxonomic assignation (at the best possible level). RF scores correspond to the mean decrease of GINI coefficient (higher scores mean a better capacity to discriminate groups).

#### 6.2 Objective #2: Understand the interaction between humic DOC and microbial consortia

We first characterized the overall DOC quality in the sampled environments using Principal Components Analysis (PCA) (Fig. 4a), PARAFAC model components (Fig. 4b) and FEEM scans (Suppl. Fig. 5 in Suppl. Mat.). Then, to better understand the interaction between humic DOC and bacterioplankton communities, we assessed to what extent humic DOC drives community assemblies and transcription profiles. We fitted the humic DOC concentration on NMDS plots (Fig. 5a,b) based on unweighted Unifrac distances using *ordisurf* [40]. Then, we assessed the Spearman correlations between the humic DOC concentration and the abundance or transcription of bacterial ASVs (Fig. 5c,d). Correlations > 0.5 with p-value < 0.05 (after Bonferroni correction) were plotted on Cytoscape v. 3.7.1. We also aimed to identify the potential pathways of lignin degradation present in our dataset. To do so, we performed a comprehensive literature search to list the potential pathways/enzymes currently known to be involved in lignin degradation in bacteria (see list of enzymes in Suppl. Table 4). We searched for the presence of these enzymes in our dataset using our functional database. When these enzymes were observed to correlate with the concentration of humic DOC, we plotted their abundance and the abundance of the taxa in which they were found using a heatmap (Fig. 6). Finally, based on our results concerning the taxon *Polynucleobacter sinensis*, we investigated whether the presence of the genes coding the enzymes from Suppl. Table 4 would be found in its reference genome [43] (SRA accession #: PRJNA295472). We processed the reference bacterial genome as previously described for the reference metagenome, to obtain a list of all pathways/enzymes inventoried.

**Figure 4:**
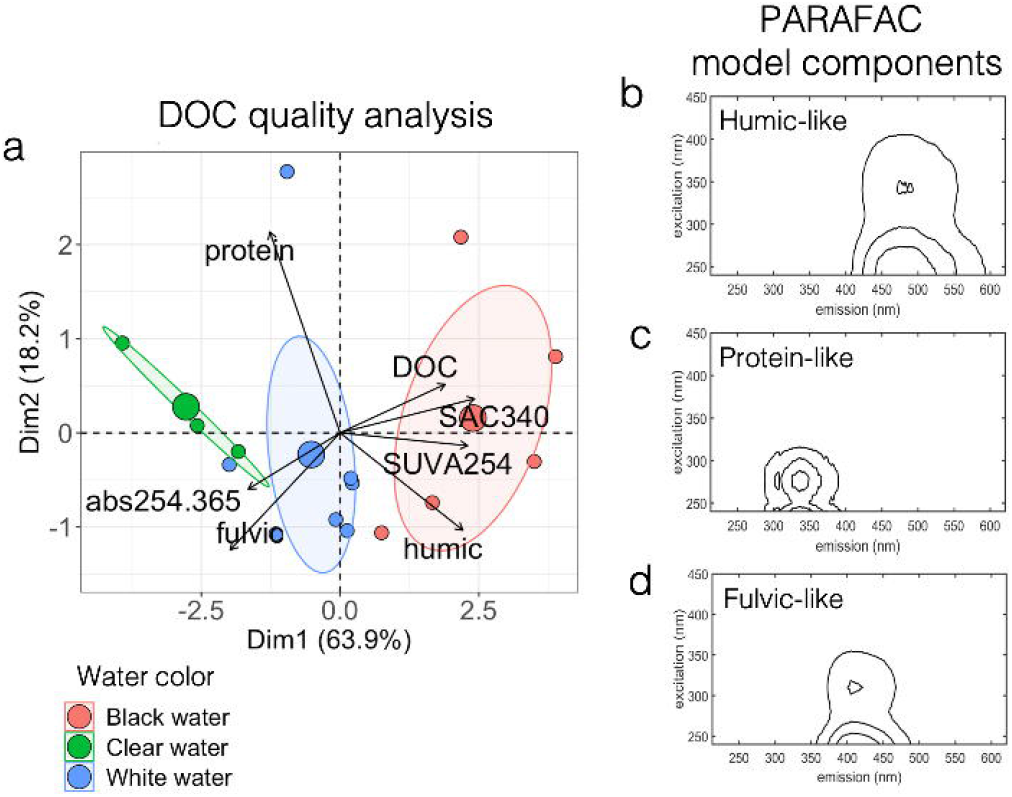
Black water sites showed unique dissolved organic carbon (DOC) quality profiles, characterized by a higher load of aromatic, high molecular weight humic carbon. (a) Principal Components Analysis showing how sampling sites cluster according to their DOC quality profiles. The three bigger dots refer to group centroids. Environmental variables: “DOC” refers to concentration (mg L^−1^) of DOC; “protein” refers to concentration of protein-like DOC; “humic” refers to concentration of humic-like DOC; “fulvic” refers to concentration of fulvic-like DOC; SAC340, SUVA254 and abs254.365 refer to absorbance ratios detailed in Table 1. Then, PARAFAC model components showed the presence of humic-like (b), protein-like (c), and fulvic-like (d) DOC fractions in the sites sampled.

**Figure 5:**
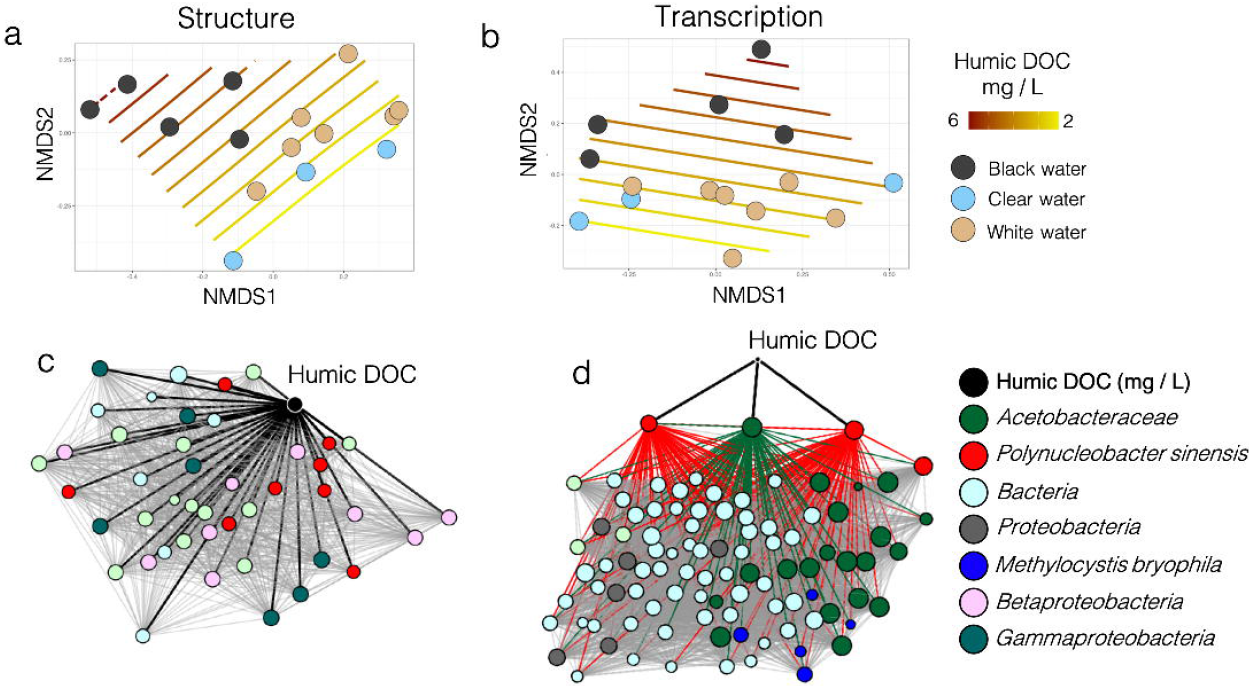
The concentration of humic dissolved organic carbon (DOC) correlated to bacterioplankton structural and transcriptional profiles’ clustering and feature abundance. Ordisurf plots (i.e. NMDS fitted with isolines describing humic DOC concentration variations) for the structural (a) and transcriptional (b) profiles. Samples were merged per sampling site and were colored according to their water type. Spearman correlation-based networks of co-abundance between humic DOC concentrations and bacterial taxa abundance (c) or transcriptional activity (d). For ease of viewing, only the taxa which correlated directly with humic DOC, or that were direct neighbors of such taxa, were kept for network construction. Nodes were colored according to their taxonomic assignation (at the best level possible). In (c,d) black edges represented direct correlations between humic DOC and specific taxa. In (d), edges from the three nodes directly correlated to humic DOC were colored according to their node taxonomic assignation to highlight the indirect effect of humic DOC on the transcriptional profile. These edges were not colored in (c) for ease of viewing.

**Figure 6:**
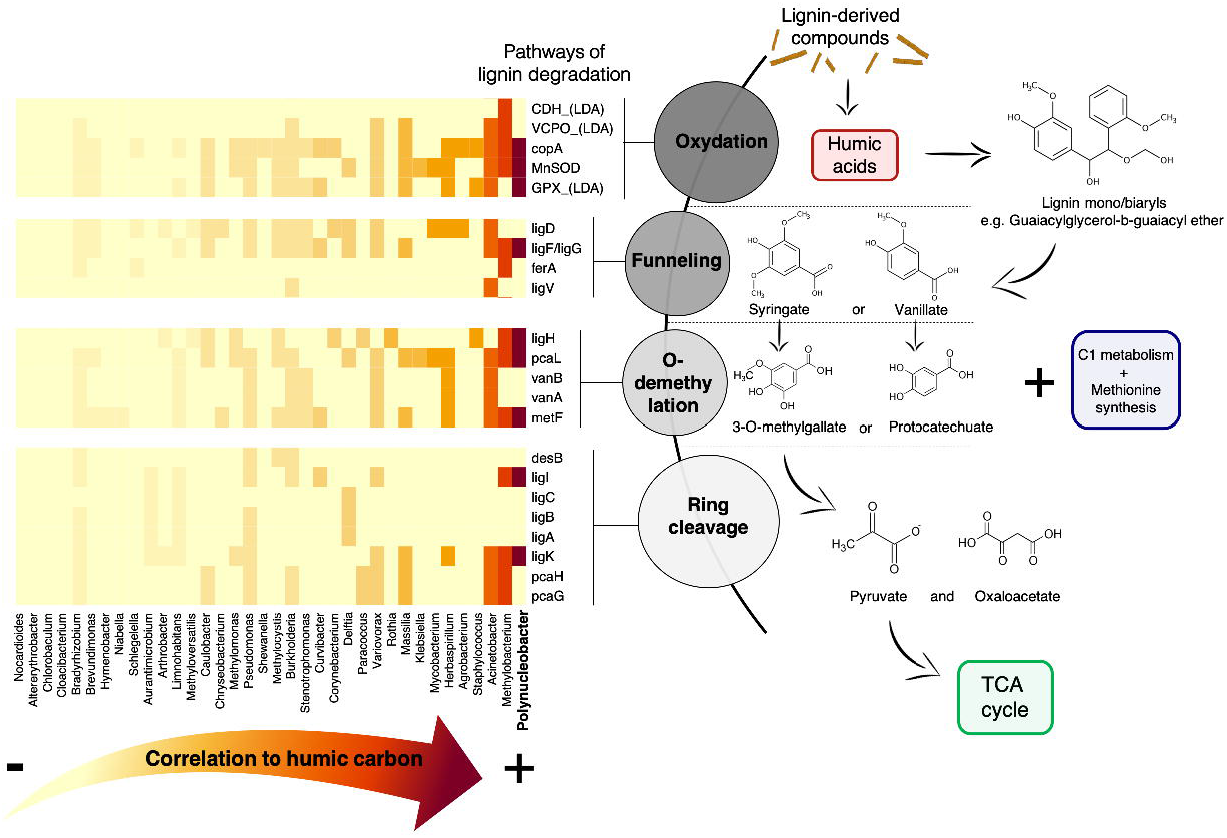
Potential presence of different pathways of the four main steps of lignin degradation (Kamimura *et al*. 2017) in the genera which abundance correlated with humic DOC concentration. Spearman correlation between humic DOC and genera abundances were plotted on a heatmap from less to most correlated (left to right respectively). *Polynucleobacter* is in bold as it is the genus showing the highest correlation to humic carbon concentration. The complete list of pathways that were screened for in the functional repertory can be found in Suppl. Table 4, however, only the pathways that were found in at least one genus are represented in the heatmap for ease of viewing. TCA cycle refers to the tricarboxylic acid cycle.

## Results

### Objective #1: Identify the environmental variables shaping the structural, transcriptional and functional profiles of Amazonian bacterioplankton

Bacterioplankton communities significantly clustered according to water type for structural, transcriptional and functional NMDS analyzes (Fig. 2a,b,c). In all cases, PERMANOVA analyzes consistently showed that water type significantly drove community structure (F = 8.5, df.res = 82, R^2^ = 0.17, p < 0.001), transcription profile (F = 2.9, df.res = 82, R^2^ = 0.06, p < 0.001), and functional repertory (F = 11.8, df.res = 82, R^2^ = 0.22, p < 0.001). NMDS plots from Fig. 2a,b,c also suggest that white and clear water communities were similar, and differed notably from black water communities which form an independent cluster. The top 10 parameters that influenced the structure, transcriptional profile and functional repertory were not identical. However, they always comprised at least one parameter associated with DOC quantity/quality, as well as several parameters associated to the environmental ionic profile (i.e. conductivity or the concentration of specific free ions).

We succeeded in implementing a machine-learning Random Forest algorithm to identify which taxa best discriminated black versus white/clear waters. The out-of-bag error rate (node error rates in the trees of classification) was only 2.35% when considering community structure, and it was 18.82% when considering the transcriptional profile. The taxa that best discriminated water types were associated to either black or white/clear waters. Strength of association to water type was stronger for taxa associated to black water than to white/clear waters, especially when looking at the transcriptional profile (Fig. 3b). This was also true for functions in the functional repertory (Suppl. Fig. 4 in Suppl. mat.). Detailed description of the ASV clusters discriminating water types can be found in Suppl. mat. Taxa showing the strongest decrease of GINI coefficient (i.e. the best association score, or discriminating power) at the structural level were two species of *Gammaproteobacteria*, which were both associated to black waters. At the transcriptional level, the strongest signal came from a group of *Polynucleobacter sinensis*, also strongly associated to black waters.

### Objective #2: Understand the interaction between humic DOC and microbial consortia

The DOC quality varied significantly within the different water types of the Amazon Basin (Fig. 4a). The PARAFAC model showed that the DOC was composed of three main fractions: humic-like, fulvic-like and protein-like fractions (Fig. 4b,c,d). Black water sites contained higher amounts of DOC, enriched in humic-like DOC of a greater aromaticity and molecular weight. White water sites contained DOC characterized by a high content of fulvic-like components, while clear water sites contained more protein-like DOC of low aromaticity and molecular weight (see Table 1 for raw data, Fig. 4 and FEEM scans on Suppl. Fig. 5 in Suppl. mat.).

The humic-like fraction of DOC was associated to the bacterioplankton community composition and transcription profiles within the different waters of the Amazon basin, as the humic DOC concentration isolines respect the natural clustering of the communities on the unweighted Unifrac NMDS plots from Fig 5a,b. A co-abundance Spearman correlation network-based approach enabled us to identify which taxa correlated to humic DOC (Fig. 5c,d). The results at the structure and transcriptional levels show different correlation profiles. Indeed, at the structural level, the results suggested a direct correlation between humic DOC concentration and 44 taxa. In contrast, at the transcriptional level, the influence seemed to be mostly indirect since there was only 3 taxa directly correlated to humic DOC concentration. However, these three taxa (two *Polynucleobacter sinensis* and one *Acetobacteraceae*) were key in the overall community transcription profile - they were important interaction hubs as their activity was strongly correlated to 93 other taxa.

We recomputed the Spearman correlation analysis (between humic DOC and bacterial taxa) after agglomerating all ASVs at the genus level, to ensure compatibility with the metagenome reference database. Several taxa that were significantly associated to humic DOC at the genus level were also associated with humic DOC at the ASV level (e.g. *Polynucleboacter* and *Methylocystis*). We investigated the presence/absence of the enzymes known to be part of lignin degradation pathways (see Suppl. Table 4) in the subset of all genera which abundance correlated to humic DOC (Fig. 6). Three genera were highly correlated with humic DOC: *Polynucleobacter*, *Methylobacterium*, and *Acinetobacter*. These three genera also possessed enzymes involved in the four main steps of lignin degradation.

Overall, the taxa *Polynucleobacter sinensis* has shown the most promising results concerning its potential implication in the degradation of lignin-derived compounds/humic DOC. Indeed, the Random-Forest analysis has shown that *P. sinensis* is the best taxa to discriminate black from white/clear waters at the transcriptional level (Fig. 3b). *P. sinensis* ASVs abundance was correlated to humic DOC concentration in structural and functional profiles (where they represented two out of three ASVs correlated to humic DOC) (Fig. 5c,d). When the correlation analysis was performed at the genus level, *Polynucleobacter* was the taxa which abundance showed the strongest correlation to humic DOC (Fig. 6). Finally, *Polynucleobacter* was involved in the four main lignin degradation steps. Thus, we investigated whether the presence of the genes coding the enzymes from Suppl. Table 4 would be found in the *P. sinensis* reference genome published by Hanh *et al.* (2009) [43]. We found that *P. sinensis* actually possesses the lignin-degradation enzymes that were found in the reference metagenome, which confirmed our result from Fig. 6 at the genus level. *P. sinensis* possesses genes coding for enzymes involved in the four steps of lignin degradation: initial oxidation, funneling, O-demethylation, and ring cleavage pathways. In brief, the set of genes found in *P. sinensis* genome suggests that it uses a derivative of the β-aryl ether degradation pathway for diaryl lignin residues, which concludes in an extradiol 4,5-PCA ring meta cleavage producing pyruvate and oxaloacetate. Further details and inference of possible degradation pathways in *P. sinensis*, as well as details on the presence of lignin degradation pathways in non bacterial sequences from the metagenome database can be found in Suppl. mat.

## Discussion

### Water type shapes bacterial community composition, transcriptional activity, and functional potential

Water type has been shown to be a major driver of the diversity, composition and population genomics of eukaryotic biological communities in Amazonia. This has been shown in a vast array of species including teleosts [44–47], phyto and zooplankton [48], as well as periphyton communities [49]. There has also been a few studies focused on Amazonian bacterioplankton, although most of them did not sample in different water types [20, 35– 38, 50–54]. The aforementioned studies were mostly focused on the dynamics of bacterial communities in the plume downstream of the Amazonian river system [38, 50, 51], or included a very limited number of sampling sites (e.g. only 1 site in [52]), or solely relied on taxonomic approaches, without considering the transcriptional activity of taxa involved [35– 37, 52–54]. In our study, we used an unprecedented approach combining data from the structure, transcription activity and functional repertory of 15 bacterioplankton communities from the three water types of the upper Amazonian basin. We were also the first to characterize extensively the environments sampled (34 parameters measured, from physicochemical parameters to primary productivity, ionic profiles, detailed DOC quality analyses, etc.) which enabled us to identify accurately which variables were most likely involved in shaping Amazonian bacterioplankton.

Our results showed an important influence of water type on bacterioplankton structure, transcription and functional profiles (Fig. 2). For all these profiles (Fig. 2a,b,c), we showed that among the 10 environmental parameters that were the most associated with community clustering, several are known to be the main parameters driving differences between water types. For instance, higher concentrations of total DOC, humic DOC, Al and Pb are known to be strongly associated with black waters [16, 55], while higher primary productivity (chlorophyll a), conductivity, and pH are characteristic of white and clear water environments. Interestingly, we observed that the clustering of functional repertory according to water types (Fig. 2c) was not as clear as structural or transcriptional profiles clusterings (Fig. 2a,b). This result might be associated to the fact that several house-keeping genes are shared by all bacterial members, thus reducing inter-site variability when considering the functional repertories. However, based on the significant PERMANOVA results (Fig. 3c) it still appears that there were significant water-type-specific functional profiles. Additionally, the Random-Forest analysis also identified 40 functions which have strong predictive capability in a black versus white/clear water model (Suppl. Fig. 4 in Suppl. mat.).

An aspect that should be considered when analyzing the functional profiles from a tailored metagenomic database is the limitation of inferring functional repertoires from 16S rRNA sequence based ASVs, in comparison to using a shotgun metagenomic approach for each bacterioplankton sample [56, 57]. Although our approach is analog to current functional inference pipelines such as PiCRUST [57], it palliates to an important caveat of these pipelines concerning the fact that they were designed primarily for human microbiome research, and thus lack the depth needed for thorough and precise characterization of aquatic bacterioplankton functions [56]. However, the main downfall of any approach using taxonomy to infer functions is that the completeness of the inferred functional profiles relies on the accuracy of taxonomical assignations, which are not always fully resolved in understudied environments like the Amazonian basin [20].

An interesting result emerging from the Random-Forest analysis (Fig. 3 and Suppl. Fig. 4) is the fact that the strength of association of taxa or functions to black water environments was stronger and more consistent than associations to the white/clear water environments. Indeed, the taxa/functions that best discriminated water types were mostly found to be associated to black water environments, which suggests that black water communities have transcriptional activities and functional repertories that are more specific than white/clear water sites which showed higher inter-site variability. This result concords with the fact that black water is an “extreme” environment offering challenging physicochemical conditions for biological processes, i.e. acidic pH (2.8-5.0) and very low conductivity (≈ 10 μS) [11]. Adaptation to these conditions potentially necessitates the presence and transcription of certain key functions, which constitute a unique signature/profile of these communities due to their omnipresence in these environments. Several studies have focused on how microorganisms populating extreme environments cope with stress [58, 59]. Most have found that genome plasticity, including codon bias, nucleotide skew and horizontal gene transfers (HGTs), enable evolutionary adaptation to extreme conditions [60–62]. For most bacteria, adaptation to extreme environments is a highly dynamic and complex process that involves multiple evolutionary forces [63, 64]. Although not tested in this study, these forces could potentially drive the appearance of a consistent signature/profile found in the microbiomes from similar hostile environments such as black waters from different sites.

### Humic DOC and bacterioplankton communities

Dissolved organic carbon forms the very basis of the majority of aquatic food webs and is an important food source to heterotrophs within river systems [65, 66]. The bioavailability of DOC to bacterioplankton depends on the type of DOC present. It has been suggested that allochthonous humic-like DOC may be more bioavailable to bacteria than lower molecular weight DOC [67] and that some bacteria prefer terrestrially derived DOC over autochthonous protein-like DOC derived from algae and/or bacteria (Roiha *et al*. 2016). Previous research has shown that bacteria are able to breakdown humic DOC, supporting the idea that this component is bioavailable to some bacterial species [67, 68]. Our analysis of DOC quality from the 15 sites showed that black waters have distinct DOC profiles, with higher concentrations of DOC, comprising a significant enrichment in the humic fraction characterized by higher SAC340 and SUVA254 scores (table 1 and Fig. 4). These results support previous findings by Holland *et al*. (2018) which suggest that naturally acidic waters show a unique DOC signature when compared to circumneutral and groundwater fed systems. Multivariate correlation analyzes also suggested that the concentration of humic DOC is an important factor shaping the structure and transcriptional activity (Fig. 2 and 5) of Amazonian bacterioplankton. Furthermore, analyzes at the functional level showed that there are several taxa in the Amazonian bacterioplankton community which abundance correlates with humic DOC, and which possess genes associated with lignin degradation processes. Overall, the taxa *Polynucleobacter sinensis* has shown the most promising results concerning its potential implication in the degradation of lignin-derived compounds within humic DOC. Indeed, the Random-Forest analysis showed that *P. sinensis* was the best taxa to discriminate black from white/clear waters at the transcriptional level (Fig. 3b). *Polynucleobacter sinensis* ASVs abundance and activity were also strongly correlated to humic DOC concentration (Fig. 5c,d).

The genus *Polynucleobacter* mostly comprises free-living aquatic bacteria and has been reported in numerous freshwater lakes and ponds [69]. Several studies have detected a strong correlation between the abundance of this genus and DOC concentration [69–71]. Some of these reports suggest that *Polynuclebacter* can feed on dissolved organic matter [70–71]. However, up to now the implication of this clade in humic carbon degradation in Amazonia has yet to be investigated. Here, we searched the reference genome of *P. sinensis* [43], the taxa found to correlate most reliably with humic DOC concentration, for genes coding for enzymes known to play key roles in the lignin degradation process. Interestingly, we report for the first time that the genome of *P. sinensis* contains genes for key enzymes from the four steps of the lignin degradation process, i.e. initial oxidation, funneling, O-demethylation, and ring cleavage pathways. However, since not all enzymes required for each pathway were detected in its genome, further studies are needed to decipher exactly if and how this bacterium degrades lignin, and to determine whether it is able to perform this process alone, or with associations (e.g. via syntrophic interactions) with other members of the bacterioplankton community, or with non-bacterial microbes. For instance, in the metagenomic reference database used for our study, we detected the presence of the gene ligD in several bacterial species but not in *P. sinensis* (Fig. 6). This gene was also detected in two fungi (*Aureobasidium* and *Ascomycota*). This enzyme catalyzes the oxidation of guaiacylglycerol-b-guaiacyl ether, one of the first steps of the β-aryl ether lignin degradation pathway for which *P. sinensis* possess several other enzymes. Although not tested here, functional compartmentalization of the community for lignin degradation has already been documented [72], and could potentially involve inter-kingdom relationships via alternative pathways that remain to be discovered (reviewed in [73, 74]). Finally, further studies should also assess if bacterial DOC degradation in Amazonia is dependent on a coupling with physical photodegradation processes, as suggested by Nalven *et al*. (2020) [75].

## Conclusion

Our study provides the first insights into the factors shaping the taxonomical, transcriptional and functional profiles of Amazonian bacterioplankton communities. Our comprehensive approach combined observations from bacterioplankton samples collected from 15 sites across the Amazon basin, to a meta-analysis of 90 Amazonian shotgun metagenomes used to build a functional reference database. The results show that the most important factors affecting these bacterioplankton communities within the three different water types of the Amazon basin are DOC, humic DOC, Pb and Al concentrations, which are all associated to black water environments. We also used a novel Random-Forest approach to highlight the taxa mostly associated to the observed differences between water types. Among these taxa, ASVs assigned to the species *Polynucleobacter sinensis* particularly stood out, as their abundance and their transcriptional activity were strongly correlated to black water environments, and specially to humic DOC concentration. Screening the reference genome of this bacteria, we found genes coding for enzymes implicated in all the main lignin degradation steps, which suggest that this bacteria may play a significant role in this important process key to carbon cycling in the Amazonian basin.

## Acknowledgements

We thank the National Geographic Society, NSERC, MITACS, and Ressources Aquatiques Québec for awarding travel grants to FÉS. This study was part of the NSERC Discovery grant of ND, the INCT ADAPTA project of ALV, and supported by a Canada-Brazil Awards – Joint Research Project of ND and ALV, by CNPq, FAPEAM and CAPES. We thank Thiago Nascimento, Reginaldo Oliveira and Nazaré Paula for technical support with field work logisitcs. We thank Roxanne Dhommée for support in the molecular biology laboratory work. Finally, we thank the anonymous reviewers that generously took the time to help improve this manuscript.

