## Supplementary Material 1 for "The Amazon River microbiome, a story of humic carbon"

**Measure of environmental parameters**

**Primary production**. Three replicates of 250 mL of water per site were filtered using a Masterflex Easy-Load II peristaltic pump from Cole-Parmer (cat #HV-77200-62, Montreal (QC), Canada) through 0.45 µm-pore size glass fiber filters, which were then immediately stored at -80°C for chlorophyll a quantification [1]. Chlorophyll a was extracted after a 24h incubation of filters in acetone at -20°C before measuring the absorbance on a Turner Designs (San Jose (CA), USA) fluorimeter model 10AU (cat #1100-100). Chlorophyll and phaeopigment concentrations were then calculated according to the method in [1, 2].

**Nutrients**. For nutrients (NO_2_^-^, NO_3_^-^, silicates) analysis, three replicates per site of 12 mL of water were filtered through 0.45 µm-pore size glass fiber filters and stored in sterile and acid-washed (HCl 1M) 15 mL Falcon® tubes (Fisher Scientific, Hampton (NH), USA). 24 µL of HgCl_2_ solution (1g/100mL) were added for conservation, before measurement on Bran and Luebbe III (AA3) nutrient autoanalyzer as described in [3].

**Dissolved organic carbon and dissolved metals**: For analysis of dissolved metals and DOC, we filtered samples through Millipore PVDF 0.45 mm Sartorius filters (Sartorius, Germany). Metal suites were measured by inductively coupled plasma mass spectroscopy (ICP-MS). Quality assurance and quality controls (QA/QC) consisted of analysis of method blanks, laboratory duplicates, and matrix spikes were conducted using certified standards. Total suspended solids was determined using gravimetric analysis as outlined in the EPA Method 160.2. DOC was analysed as non purgeable organic carbon using a total carbon analyzer (Apollo 9000 combustion TOC analyzer: ©Teledyne Tekmar, Mason, USA). The TOC machine was calibrated using primary standard grade potassium hydrogen phthalate (KHP) and QA/QC KHP standards were ran every 10 samples. Fluorescence excitation emission (FEEM) and absorbance scans were performed using a quartz cuvette in an Aqualog® fluorimeter (cat# Aqualog, HORIBA Scientific, Piscataway (NJ), USA) to determine DOC components and characteristics. FEEM scans along with simultaneous absorbance measurements were conducted on all samples, with excitation wavelengths in 2-nm steps between 250-450 nm, and emission wavelengths of 250-620 nm. The absorbance of a blank of ultrapure water was run before each sample and automatically subtracted from each sample, with inner filter effects and 1^st^ and 2^nd^ order Rayleigh and Raman scatter also removed. The FEEMs were analyzed in MATLAB R2014b (MathWorks, Inc. © 1994-2016) and modelled using parallel factor analysis (PARAFAC) (PLS-toolbox in MATLAB: Eigenvectors Research Inc, WA, USA). The PARAFAC model was validated following the recommendations in [4]. No clear consistent patterns and peaks were visible in the residual plots, core consistency was 99%, and split-half analysis results were consistent with the model. Relative DOC aromaticity and molecular weight were determined using specific UV absorbance at 254nm (SUVA_254_), and at 350nm (SAC_340_) along with the absorbance ratio at 254nm and 365 (abs_254/365_).

**Ionic composition**: Water samples for determination of ionic composition (Na^+^, Ca^2+^, Mg^2+^, K^+^) were analysed using flame atomic absorption spectroscopy (Perkin-Elmer model 3100, cat #63929-1, Perkin-Elmer Inc, Woodbridge (ON), Canada). Cl^-^ was measured using the colorimetric method described by [5]. Hardness was calculated from the Ca^2+^ and Mg^2+^ concentrations.

**16S sequence processing**

In brief, quality control of reads was done with the filterAndTrim function using the following parameters: 290 for the forward read truncation length, 270 for the reverse read truncation length, 2 as the phred score threshold for total read removal, and a maximum expected error of 2 for forward reads and 3 for reverse reads. The filtered reads were then fed to the error rate learning, dereplication, and ASV inference steps using the functions learnErrors, derepFastq, and DADA, which are all from the DADA2 pipeline [6]. Chimeric sequences were removed using the removeBimeraDenovo function with the “consensus” method parameter. Sequenced PCR negative controls were used to remove ASVs identified as potential cross contaminants using the isContaminant function from the “decontam” package in R with a threshold of 0.4. Taxonomic annotation of amplicon sequence variants (ASV) was performed by using blastn matches against NCBI “16S Microbial” database. As the NCBI database for 16S sequences is updated more frequently than other sources, it matched our requirements for exhaustive information about lesser-known taxa, while minimizing ambiguous annotations. Matches above 99% identity were assigned the reported taxonomic identity. Sequences with no matches above the identity threshold were assigned taxonomy using a lowest common ancestor method generated on the top 50 blastn matches obtained. This method is closely inspired from the LCA algorithm implemented in MEGAN [7].

**Shotgun metagenome database construction**

The following datasets were fetched from the NCBI Sequence reads archive (SRA) : SRR1182511, SRR1182512, SRR1183643, SRR1183650, SRR1185413, SRR1185414, SRR1186214, SRR1199270, SRR1199271, SRR1199272, SRR1202081, SRR1202089, SRR1202090, SRR1202091, SRR1202095, SRR1204580, SRR1204581, SRR1205250, SRR1205251, SRR1205252, SRR1205253, SRR1209976, SRR1209977, SRR1209978, SRR1514963.1, SRR1515032.1, SRR1518285.1, SRR1522964.1, SRR1522971.1, SRR1522973.1, SRR1522974.1, SRR1786279, SRR1786281, SRR1786608, SRR1786616, SRR1787940, SRR1787943, SRR1788318, SRR1790487, SRR1790489, SRR1790644, SRR1790646, SRR1790647, SRR1790676, SRR1790678, SRR1790679, SRR1790680, SRR1792674, SRR1792852, SRR1793861, SRR1793862, SRR1796116, SRR1796118, SRR1796234, SRR1796236, SRR4831644, SRR4831645, SRR4831646, SRR4831647, SRR4831648, SRR4831649, SRR4831650, SRR4831651, SRR4831652, SRR4831653, SRR4831654, SRR4831655, SRR4831656, SRR4831657, SRR4831658, SRR4831659, SRR4831660, SRR4831661, SRR4831662, SRR4831663, SRR4831664, SRR4831665, SRR4831666, SRR4831667, SRR4833053, SRR4833055, SRR4833056, SRR4833057, SRR4833059, SRR4833060, SRR4833062, SRR4833064, SRR4833067, SRR4833073, SRR4833077, SRR4833080, SRR4833081, SRR4833084, SRR4833086, SRR4833087, SRR4833089, SRR5123271, SRR5123272, SRR5123273, SRR5123274, SRR5123275, SRR5123276 and SRR5123277. These reads were pooled into forward and reverse reads (when possible) and assembled using the megahit assembler with the large metagenome (meta-large) parameter preset, a k-min of 27 and with iterative increments of k (k-step) of 10. Taxonomic annotation of the assembled contigs was performed by using blastn matches against the NCBI “16S Microbial” database. Sequences with no matches above the identity threshold were assigned taxonomy using a lowest common ancestor method generated on the top 50 blastn matches obtained. Functional annotation of contigs was made using the following steps: First, predicted proteins were determined from the contig nucleotide sequences using ORFM [8]. Then, predicted protein sequences were annotated using blastp against the swissprot-uniprot database in order to obtain gene ontology (GO) and KEGG ontology (KO) information. Finally, a database combining the taxonomic and functional information was made to be used as reference for subsequent steps.

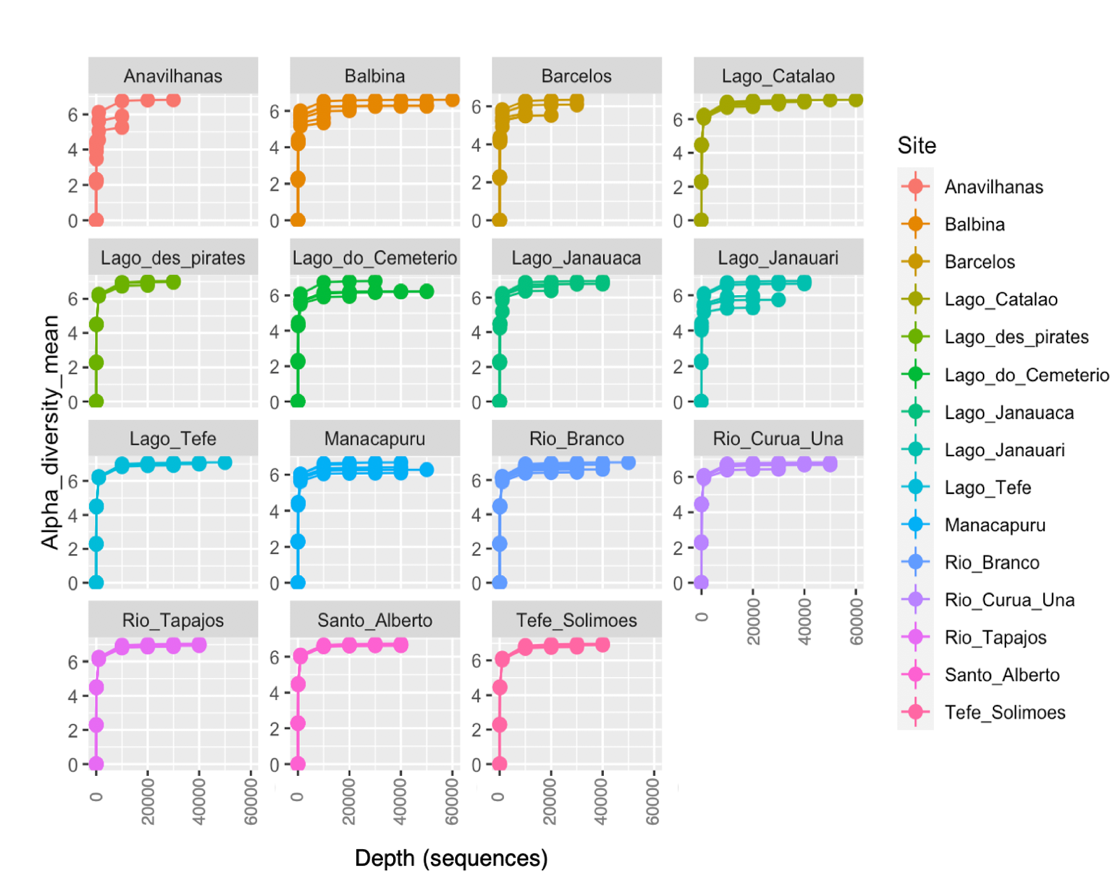

**Suppl. Figure 1**: Rarefaction plots of the samples for each sampling site, for the **structural** analysis. The rarefaction analysis was based on the Shannon diversity for each sample group, according to the sequencing depth (number of sequences used).

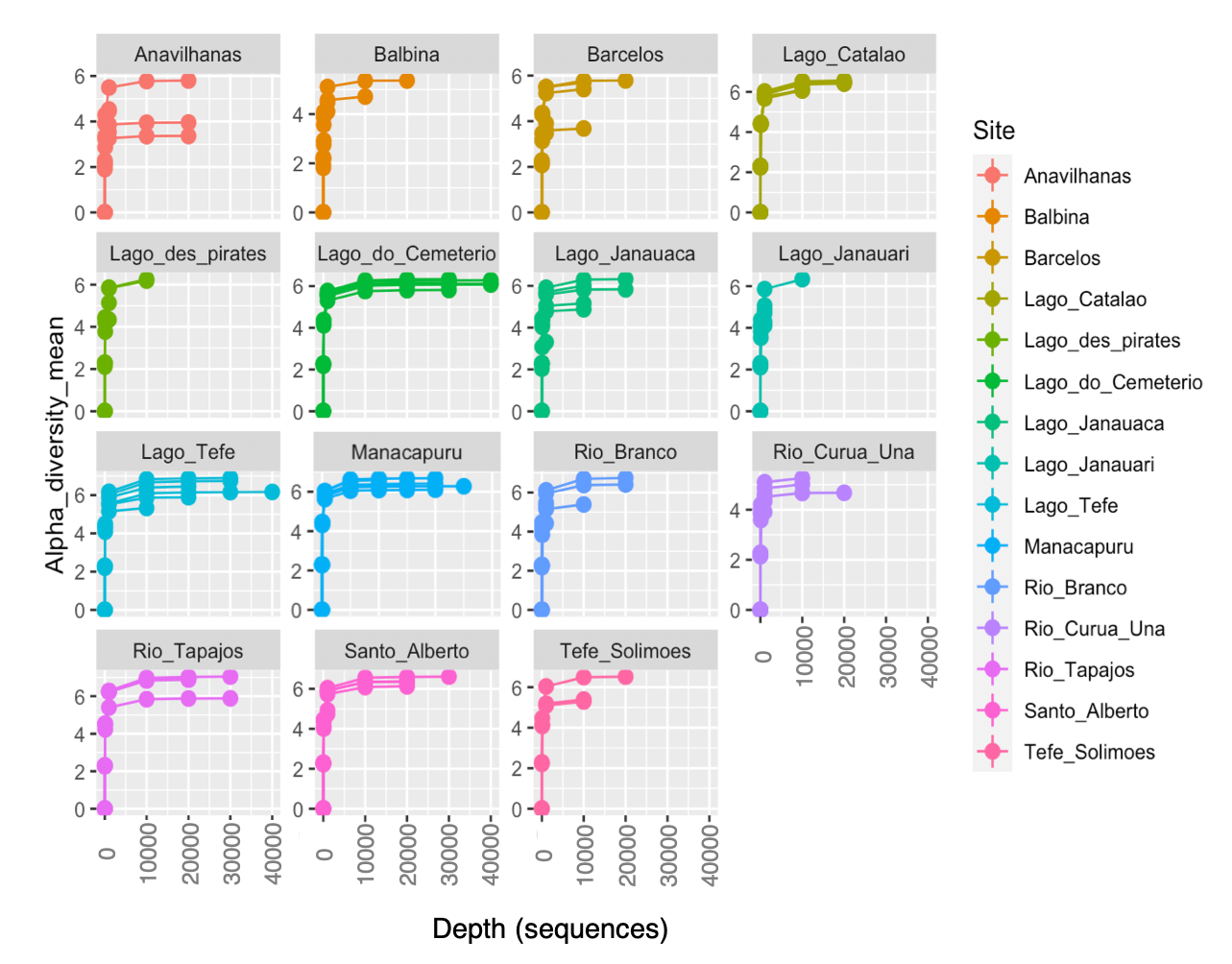

**Suppl. Figure 2**: Rarefaction plots of the samples for each sampling site, for the **transcriptional activity** analysis. The rarefaction analysis was based on the Shannon diversity for each sample group, according to the sequencing depth (number of sequences used)

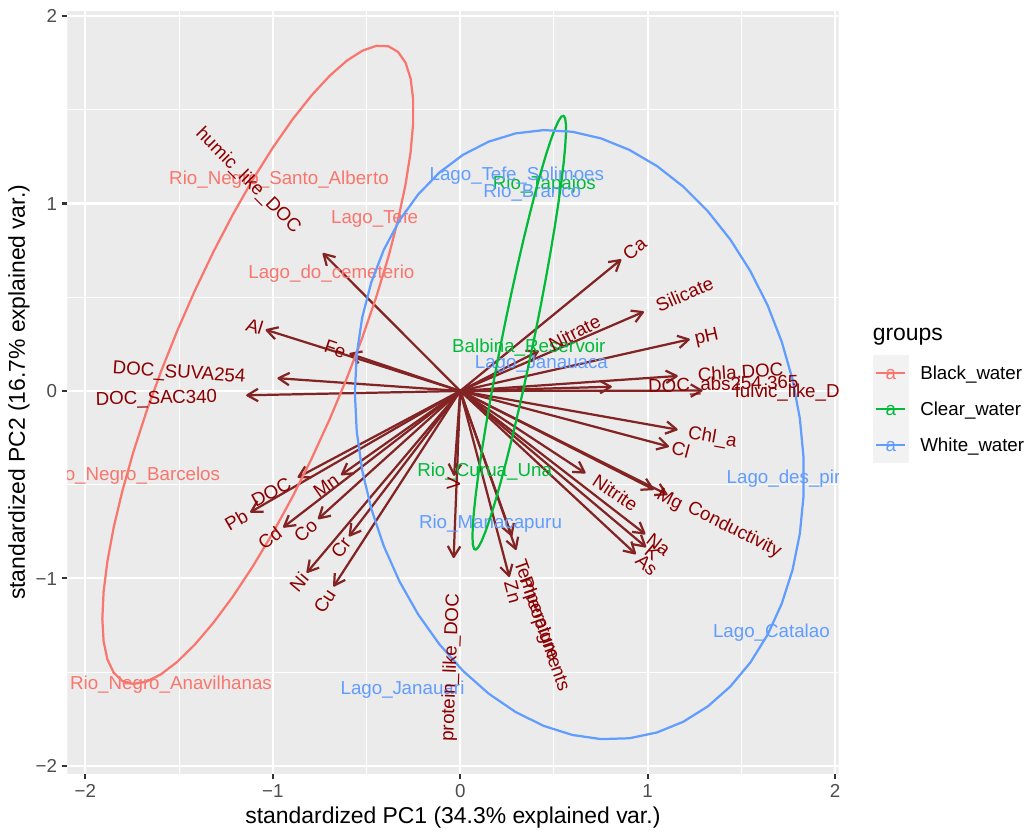

**Suppl. Figure 3**: Principal Components Analysis of the water parameters measured at each sampling site. The color of the label of each site corresponds to the water type found at that site.

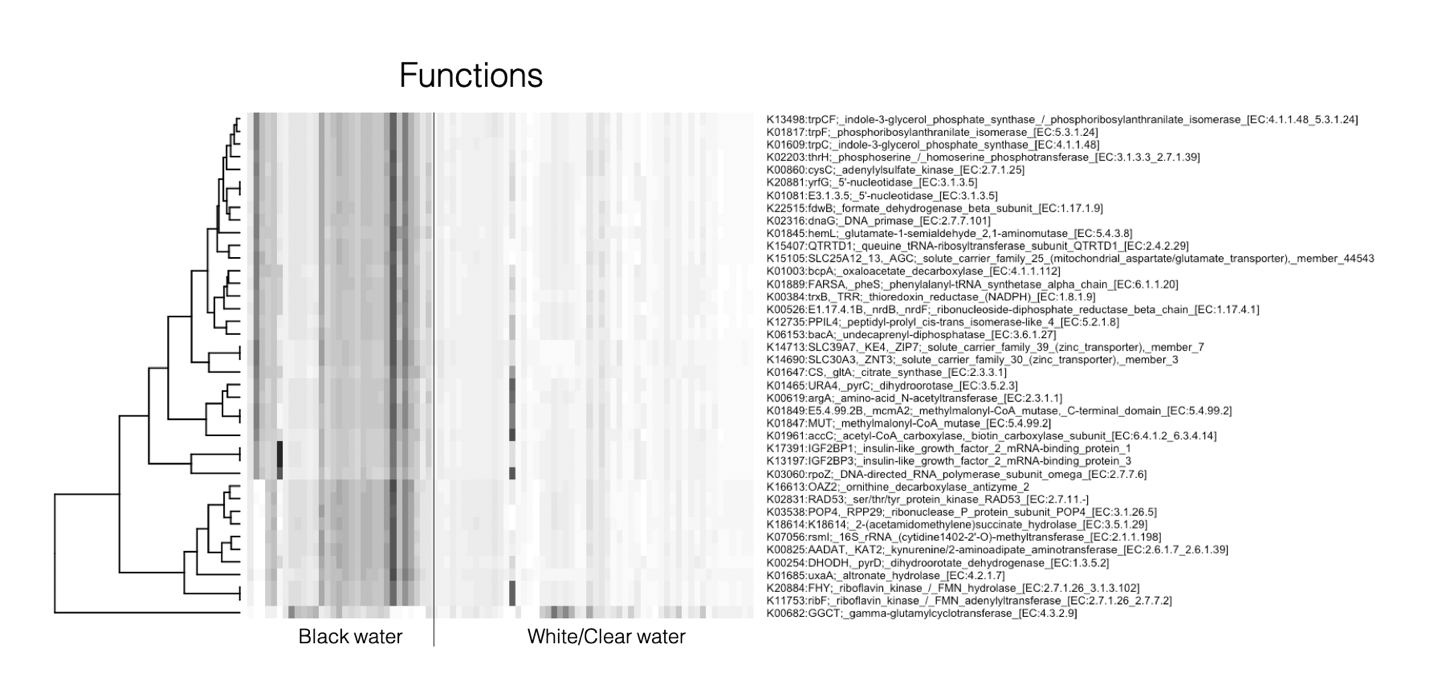

**Suppl. Figure 4**: Random-Forest (RF) machine-learning analysis identifies the 40 functions showing the most important differentiation between black and white/clear water colors. Heatmap columns represent samples and rows are different functions.

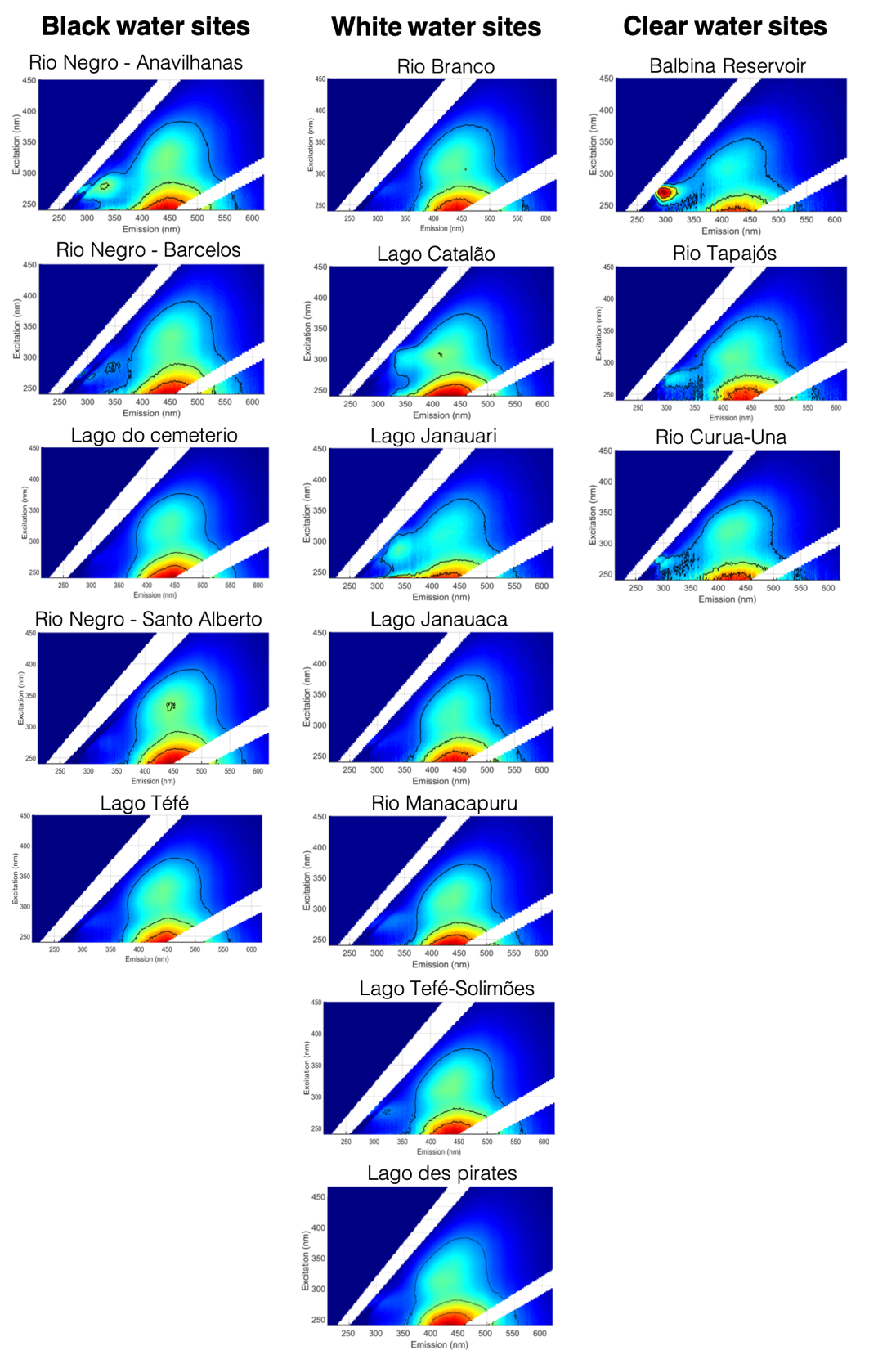

**Suppl. Figure 5**: Fluorescence Excitation Emission Scans (FEEMs) of the DOC from all sites.

**Results: ASV clusters discriminating water types (Fig. 3)**

At the community structure level, ASVs which best discriminated black versus white/clear waters included two clusters for each water type (Fig. 3a). One cluster associated to white/clear comprised only 11 *Actinobacteria*, while the other comprised seven *Actinobacteria*, one *Alphaproteobacteria* and one *Burkholderiales*. One of the two clusters associated to black water comprised two *Polynucleobacter sinensis* and one *Acetobacteraceae*, while the other one, more diverse, comprised seven *Gammaproteobacteria*, three unidentified bacteria, one *Acetobacteraceae* and six *Actinobacteria*. At the transcriptional level, we found only one cluster associated to each water type (Fig. 3b). The cluster associated to black water comprised 10 *Acetobacteraceae*, 10 *Polynucleobacter sinensis* and one *Actinobacteria*, while the one associated to white/clear waters was more diversified and comprised six Actinobacteria, two *Geobacillus stearothermophilus*, one *Caulobacteraceae*, two *Delftia acidovorans*, three *Bradyrhizobium namibiense* and four unidentified bacteria. Two taxa that were always associated to black water environments (at the structure and transcriptional levels) (i.e. *Polynucleobacter sinensis* and *Acetobacteraceae*), while no such consistent association was found for white/clear water environments.

**Results: Pathways of lignin degradation in *Polynucleobacter sinensis***

**Lignin oxidation**: *P. sinensis* possesses multicopper oxidases (copA, K17686) and manganese-dependant superoxide dismutases (SOD1, K04565) known to oxidise lignin derived compounds. Moreover, *P. sinensis* also has certain lignin degrading auxiliary enzymes (unable to oxidise lignin on their own, but essential to the process) such as glutathione peroxidases (GPX, K00432). **Funneling**: in *P. sinensis* oxidised lignin then seems to follow a derivative of the β-aryl ether degradation pathway for diaryl residues. Indeed, *P. sinensis* contains the glutathione S-transferases ligF/ligG (GST, K00799), performing one of the main reactions of this funneling pathway leading to the production of vanillate. **O-demethylation**: it is not clear how the O-demethylation of vanillate occurs in *P. sinensis*, as the enzyme ligM (K15066) – known to be associated to this step in other bacteria [9] – was not found in the published *P. sinensis* genome. However, it appears that a similar enzyme performs this reaction since *P. sinensis* possesses enzymes associated with the metabolism of protocatechuate (PCA), the product of the ligM O-demethylation, such as 3-oxoadipate enol-lactonase/4-carboxymuconolactone decarboxylase (pcaL, K14727). The ligM reaction is tetrahydrofolate (H_4_-folate) dependant since it adds the methyl group from vanillate to H_4_-folate to produce 5-methyl-H_4_-folate. Moreover, *P. sinensis* potentially possesses a similar H_4_-folate-dependant demethylase since we found a consortium of enzymes (i.e. metF, SHMT, and fold; K00297, E.C.:2.1.2.1, K01491) associated to the conversion of 5-methyl-H_4_-folate to 5,10-methylene-H_4_-folate and 10-formyl-H_4_-folate in its genome. These compounds can be used for the synthesis of methionine, thymidine and purine, respectively, and can also be incorporated in the bacterial C1 metabolism [10]. **Ring cleavage**: finally, it appears that *P. sinensis* is performing an extradiol 4,5-PCA ring meta cleavage, producing pyruvate and oxaloacetate, as it possesses the enzymes ligI (a 2-pyrone-4,6-dicarboxylate lactonase, K10221) and ligK (a 4-hydroxy-4-methyl-2-oxoglutarate aldolase, K10218) associated with this pathway.

In addition, to address potential complementarity with bacteria-mediated degradation pathways, we also investigated the presence of lignin degradation pathways in the non-bacterial sequences from the metagenome database. We found only three potential enzymes implicated in lignin degradation in fungi: ligD (K01971), ligH (K01938), and GPX (K00432). We could not investigate this directly from our field sampling data as we relied on a 16S-based approach, thus, the abundance of these genes in different water types could not be assessed.

**Supplementary Tables**

**Suppl. Table 1**: Concentrations of free ions and nutrients.

| **Site #** | **Water color** | **Ions: mg L^-1^** | | | | | **Nutrients: umol L^-1^** | | |
| --- | --- | --- | --- | --- | --- | --- | --- | --- | --- |
|  |  | Na^+^ | Mg^+2^ | K^+^ | Ca^+2^ | Cl^-^ | Nitrite | Nitrate | Silicate |
| 1 | Black | 0.46 | 0.12 | 0.42 | 0.04 | 0.11 | 0.11 | 3.20 | 64.41 |
| 2 | Black | 0.25 | 0.09 | 0.33 | 0.49 | 1.16 | 0.10 | 2.87 | 92.32 |
| 3 | Black | 1.80 | 0.26 | 0.65 | 0.08 | 0.32 | 0.09 | 4.36 | 72.55 |
| 4 | Black | 0.23 | 0.05 | 0.14 | 0.37 | 0.64 | 0.01 | 0.47 | 76.82 |
| 5 | Black | 0.87 | 0.19 | 0.56 | 0.82 | 0.53 | 0.08 | 4.09 | 217.19 |
| 6 | White | 1.15 | 0.43 | 0.70 | 0.93 | 1.10 | 0.04 | 8.23 | 180.48 |
| 7 | White | 1.99 | 0.20 | 0.79 | 0.06 | 1.47 | 0.19 | 1.31 | 98.31 |
| 8 | White | 4.56 | 3.76 | 1.71 | 0.83 | 1.75 | 0.09 | 0.56 | 242.31 |
| 9 | White | 3.32 | 1.00 | 1.07 | 0.44 | 2.17 | 0.13 | 20.45 | 156.51 |
| 10 | White | 4.91 | 0.14 | 1.45 | 0.05 | 1.43 | 0.12 | 1.53 | 126.01 |
| 11 | White | 1.95 | 0.21 | 0.28 | 1.11 | 1.29 | 0.03 | 6.47 | 326.53 |
| 12 | White | 5.35 | 1.76 | 1.28 | 1.17 | 3.26 | 0.61 | 11.96 | 222.31 |
| 13 | Clear | 0.80 | 0.14 | 0.67 | 0.03 | 0.78 | 0.05 | 1.55 | 85.93 |
| 14 | Clear | 0.43 | 0.47 | 0.57 | 0.68 | 0.39 | 0.09 | 1.91 | 179.36 |
| 15 | Clear | 1.52 | 0.26 | 0.67 | 0.04 | 1.22 | 0.06 | 2.55 | 171.96 |

**Suppl. Table 2**: Primary productivity characterization and measure of several physicochemical parameters. “Chl a” means the concentration of chlorophyll a; “Phaeopig.” means the concentration of phaeopigments; “Chla/DOC” is a ratio of the concentration of chlorophyll a divided by the concentration of DOC; “Temp. °C” means the temperature in ° Celsius; “Cond. uS” means the conductivity in microsiemens; “% sat. O_2_” means the percentage of saturation of dissolved oxygen.

| **Site #** | **Water color** | **Primary productivity: ug L^-1^** | | | **Physicochemical parameters** | | | |
| --- | --- | --- | --- | --- | --- | --- | --- | --- |
|  |  | Chl a | Phaeopig. | Chla/DOC | Temp. °C | Cond. uS | pH | % sat. O_2_ |
| 1 | Black | 0.35 | 2.43 | 0.03 | 31.60 | 13.10 | 3.71 | 92.12 |
| 2 | Black | 0.73 | 0.33 | 0.06 | 30.60 | 10.60 | 4.16 | 58.00 |
| 3 | Black | 0.05 | 0.38 | 0.00 | 30.70 | 13.20 | 4.24 | 53.20 |
| 4 | Black | 1.35 | 1.44 | 0.14 | 32.40 | 7.20 | 3.83 | 76.90 |
| 5 | Black | 1.82 | 1.73 | 0.26 | 30.00 | 10.60 | 4.98 | 61.50 |
| 6 | White | 6.21 | 2.89 | 1.03 | 31.00 | 22.00 | 6.25 | 88.70 |
| 7 | White | 4.62 | 17.31 | 0.65 | 32.90 | 22.40 | 4.38 | 60.00 |
| 8 | White | 7.14 | 6.60 | 0.79 | 32.90 | 174.80 | 5.70 | 44.00 |
| 9 | White | 1.35 | 1.88 | 0.24 | 29.30 | 88.00 | 6.75 | 82.60 |
| 10 | White | 2.78 | 10.54 | 0.35 | 32.60 | 24.30 | 5.31 | 72.80 |
| 11 | White | 4.41 | 3.20 | 0.77 | 30.30 | 19.70 | 6.05 | 68.60 |
| 12 | White | 9.05 | 4.69 | 1.40 | 31.90 | 127.60 | 7.15 | 31.90 |
| 13 | Clear | 0.83 | 0.78 | 0.17 | 33.20 | 16.80 | 5.05 | 103.20 |
| 14 | Clear | 2.15 | 1.03 | 0.81 | 30.00 | 14.10 | 6.36 | 80.20 |
| 15 | Clear | 1.25 | 2.38 | 0.28 | 31.20 | 19.00 | 6.00 | 79.10 |

**Suppl. Table 3**: Concentration of dissolved metals in ug/L.

| **Site #** | **Water color** | **Metals (ug/l)** | | | | | | | | | | |  |
| --- | --- | --- | --- | --- | --- | --- | --- | --- | --- | --- | --- | --- | --- |
|  |  | Al | V | Cr | Mn | Fe | Co | Ni | Cu | Zn | As | Cd | Pb |
| 1 | Black | 137.75 | 0.38 | 0.30 | 7.38 | 166.63 | 0.13 | 1.93 | 10.36 | 33.48 | 0.16 | 0.09 | 1.43 |
| 2 | Black | 150.00 | 0.10 | 0.05 | 5.90 | 160.00 | 0.10 | 0.15 | 0.30 | 11.00 | 0.05 | 0.02 | 0.27 |
| 3 | Black | 36.33 | 0.34 | 0.37 | 9.24 | 142.38 | 0.28 | 3.23 | 9.25 | 72.92 | 0.48 | 0.21 | 1.11 |
| 4 | Black | 87.00 | 0.30 | 0.05 | 4.60 | 100.00 | 0.10 | 0.33 | 1.90 | 9.00 | 0.08 | 0.13 | 0.30 |
| 5 | Black | 62.00 | 0.10 | 0.33 | 13.00 | 220.00 | 0.10 | 0.52 | 0.60 | 4.40 | 0.19 | 0.02 | 0.12 |
| 6 | White | 38.00 | 0.20 | 0.05 | 0.51 | 230.00 | 0.10 | 0.14 | 0.80 | 2.60 | 0.07 | 0.02 | 0.26 |
| 7 | White | 65.50 | 0.78 | 0.40 | 9.85 | 269.28 | 0.10 | 0.85 | 16.19 | 44.15 | 0.47 | 0.06 | 0.67 |
| 8 | White | 1.81 | 0.17 | 0.10 | 0.61 | 5.84 | 0.10 | 0.48 | 2.20 | 171.78 | 0.99 | 0.02 | 0.03 |
| 9 | White | 28.02 | 1.45 | 0.09 | 11.25 | 166.97 | 0.10 | 0.58 | 2.73 | 1.83 | 0.70 | 0.03 | 0.25 |
| 10 | White | 13.47 | 0.85 | 0.21 | 4.64 | 97.85 | 0.10 | 1.12 | 2.11 | 25.85 | 0.38 | 0.08 | 0.16 |
| 11 | White | 49.00 | 0.30 | 0.11 | 0.68 | 250.00 | 0.10 | 0.41 | 0.50 | 2.70 | 0.27 | 0.02 | 0.21 |
| 12 | White | 27.00 | 0.20 | 0.06 | 4.60 | 82.00 | 0.10 | 0.60 | 1.70 | 8.10 | 1.30 | 0.02 | 0.11 |
| 13 | Clear | 10.29 | 0.05 | 0.05 | 0.23 | 16.85 | 0.10 | 0.13 | 0.56 | 4.49 | 0.14 | 0.02 | 0.05 |
| 14 | Clear | 5.00 | 0.10 | 0.05 | 0.05 | 7.00 | 0.10 | 0.10 | 0.50 | 3.70 | 0.07 | 0.02 | 0.03 |
| 15 | Clear | 18.49 | 0.17 | 0.58 | 12.31 | 52.88 | 0.11 | 1.18 | 2.12 | 23.94 | 0.64 | 0.07 | 0.25 |

**Suppl. Table 4**: List of enzymes known to play a role in bacterial lignin degradation processes from the literature.

| **Step** | **KEGG ID** | **Enzyme name** |
| --- | --- | --- |
| Oxydation | K15733 | E1.11.1.19; dye decolorizing peroxidase [EC:1.11.1.19] |
| Oxydation | K05909 | E1.10.3.2; laccase [EC:1.10.3.2] |
| Oxydation | K17686 | copA, ctpA, ATP7; P-type Cu+ transporter [EC:7.2.2.8] |
| Oxydation | K04564 | SOD2; superoxide dismutase [EC:1.15.1.1] |
| Oxydation | K04565 | SOD1; superoxide dismutase [EC:1.15.1.1] |
| Oxydation | K16627 | SOD3; superoxide dismutase [EC:1.15.1.1] |
| Oxydation | K03782 | katG; catalase-peroxidase [EC:1.11.1.21] |
| Oxydation | K23515 | LPO; lignin peroxidase [EC:1.11.1.14] |
| Oxydation | K20205 | mpn; manganese peroxidase [EC:1.11.1.13] |
| Oxydation | K20929 | GLX; glyoxal/methylglyoxal oxidase [EC:1.2.3.15] |
| Oxydation | K17990 | VCPO; vanadium chloroperoxidase [EC:1.11.1.10] |
| Oxydation | K21820 | APO1; unspecific peroxygenase [EC:1.11.2.1] |
| Oxydation | K19813 | gdh; glucose dehydrogenase [EC:1.1.5.9] |
| Oxydation | K23272 | P2OX; pyranose oxidase [EC:1.1.3.10] |
| Oxydation | K19069 | CDH; cellobiose dehydrogenase (acceptor) [EC:1.1.99.18] |
| Oxydation | K00432 | gpx, btuE, bsaA; glutathione peroxidase [EC:1.11.1.9] |
| Funneling of monoaryls | K18383 | ferB; feruloyl-CoA hydratase/lyase [EC:4.1.2.61] |
| Funneling of monoaryls | K12508 | fcs; feruloyl-CoA synthase [EC:6.2.1.34] |
| Funneling of monoaryls | K05337 | fer; ferredoxin |
| Funneling of monoaryls | K13310 | desV, eryCI; dTDP-3-amino-3,4,6-trideoxy-alpha-D-glucose transaminase [EC:2.6.1.106] |
| Funneling of monoaryls | K21802 | vdh; vanillin dehydrogenase [EC:1.2.1.67] |
| Funneling of diaryls | K15063 | ligW; 5-carboxyvanillate decarboxylase |
| Funneling of diaryls | K15060 | ligX; 5,5'-dehydrodivanillate O-demethylase |
| Funneling of diaryls | K15062 | ligY; OH-DDVA meta-cleavage compound hydrolase |
| Funneling of diaryls | K15061 | ligZ; OH-DDVA oxygenase |
| Funneling of diaryls | K00799 | GST, gst; glutathione S-transferase [EC:2.5.1.18] |
| Funneling of diaryls | K22465 | bzaA B; 5-hydroxybenzimidazole synthase [EC:4.1.99.23] |
| Funneling of diaryls | K21568 | PLR; pinoresinol/lariciresinol reductase [EC:1.23.1.1 1.23.1.2 1.23.1.3 1.23.1.4] |
| Funneling of diaryls | K01971 | ligD; bifunctional non-homologous end joining protein LigD [EC:6.5.1.1] |
| Funneling of diaryls | K01975 | thpR; RNA 2',3'-cyclic 3'-phosphodiesterase [EC:3.1.4.58] |
| O-demethylation | K00297 | metF, MTHFR; methylenetetrahydrofolate reductase (NADPH) [EC:1.5.1.20] |
| O-demethylation | K01938 | fhs; formate--tetrahydrofolate ligase [EC:6.3.4.3] |
| O-demethylation | K15066 | ligM; vanillate/3-O-methylgallate O-demethylase [EC:2.1.1.341] |
| O-demethylation | K15064 | desA; syringate O-demethylase [EC:2.1.1.-] |
| O-demethylation | K23526 | gcoA; aromatic O-demethylase, cytochrome P450 subunit [EC:1.14.14.-] |
| O-demethylation | K23527 | gcoB; aromatic O-demethylase, reductase subunit [EC:1.6.2.-] |
| O-demethylation | K03862 | vanA; vanillate monooxygenase [EC:1.14.13.82] |
| O-demethylation | K03863 | vanB; vanillate monooxygenase ferredoxin subunit |
| O-demethylation | K14727 | pcaL; 3-oxoadipate enol-lactonase / 4-carboxymuconolactone decarboxylase [EC:3.1.1.24 4.1.1.44] |
| Ring cleavage | K10221 | ligI; 2-pyrone-4,6-dicarboxylate lactonase [EC:3.1.1.57] |
| Ring cleavage | K10218 | ligK, galC; 4-hydroxy-4-methyl-2-oxoglutarate aldolase [EC:4.1.3.17] |
| Ring cleavage | K19338 | nac; LysR family transcriptional regulator, nitrogen assimilation regulatory protein |
| Ring cleavage | K09788 | prpF; 2-methylaconitate isomerase [EC:5.3.3.-] |
| Ring cleavage | K04099 | desB, galA; gallate dioxygenase [EC:1.13.11.57] |
| Ring cleavage | K15065 | desZ; 3-O-methylgallate 3,4-dioxygenase [EC:1.13.11.-] |
| Ring cleavage | K04100 | ligA; protocatechuate 4,5-dioxygenase, alpha chain [EC:1.13.11.8] |
| Ring cleavage | K04101 | ligB; protocatechuate 4,5-dioxygenase, beta chain [EC:1.13.11.8] |
| Ring cleavage | K10219 | ligC; 2-hydroxy-4-carboxymuconate semialdehyde hemiacetal dehydrogenase [EC:1.1.1.312] |
| Ring cleavage | K10220 | ligJ; 4-oxalmesaconate hydratase [EC:4.2.1.83] |
| Ring cleavage | K00448 | pcaG; protocatechuate 3,4-dioxygenase, alpha subunit [EC:1.13.11.3] |
| Ring cleavage | K00449 | pcaH; protocatechuate 3,4-dioxygenase, beta subunit [EC:1.13.11.3] |
